# The human AMPKγ3 R225W mutation does neither enhance basal AMPKγ3-associated activity nor glycogen in human or mouse skeletal muscle

**DOI:** 10.1101/2023.08.28.555048

**Authors:** Nicolas O. Eskesen, Rasmus Kjøbsted, Jesper B. Birk, Nicolai S. Henriksen, Nicoline R. Andersen, Stine Ringholm, Henriette Pilegaard, Christian K. Pehmøller, Jørgen F. P. Wojtaszewski

## Abstract

**Background:** AMP-activated protein kinase (AMPK) is activated during cellular energy perturbation. AMPK is composed of three subunits and several variants of AMPK complexes are expressed in skeletal muscle. The regulatory AMPKγ3 subunit is predominantly expressed in fast-twitch muscle fibers. A human AMPKγ3 R225W mutation has been described in two families. In cultured cells derived from R225W carrier muscle, the mutation was reported to increase total AMPK activity. In addition, elevated glycogen levels were observed in skeletal muscle. This led to the idea of AMPKγ3 being involved in glycogen levels in skeletal muscle. Evidence for such a causative link has never been provided.

**Results:** We studied newly obtained muscle biopsies from three human carriers of the AMPKγ3 R225W mutation and matched controls and we developed a novel knock-in mouse model carrying the AMPKγ3 R225W mutation (KI HOM). In all three human pairs, the basal AMPKγ3-associated activity was reduced when assayed in the absence of exogenous AMP. No difference was observed when assayed under AMP saturation, which was supported by findings in muscle of KI HOM mice. Furthermore, the well-established stimulatory effects of AICAR/muscle contraction on AMPKγ3-associated activity were absent in muscle from KI HOM mice. Muscle glycogen levels were not affected by the mutation in human carriers or in KI HOM mice.

**Conclusions:** The AMPKγ3 R225W mutation does not impact AMPK-associated activity in mature human skeletal muscle and the mutation is not linked to glycogen accumulation. The R225W mutation ablates AMPKγ3-associated activation by AICAR/muscle contractions, presumably through loss of nucleotide binding.

## INTRODUCTION

Skeletal muscle contraction perturbs cellular energy homeostasis and initiates metabolic regulatory pathways to provide energy for the working muscle and replenish energy stores. One key signaling hub in this regulation is the AMP-activated protein kinase (AMPK). AMPK reacts to cellular energy disturbance primarily by sensing changes in adenosine nucleotide concentrations (1).

AMPK is a heterotrimeric protein complex. It consists of one catalytic (α) and two distinct regulatory subunits (β and γ). The AMPK subunits exist in several isoforms (α1/α2, β1/β2 and γ1-γ3) which are encoded by individual genes (2). The AMPK subunits are expressed in a tissue- and species-dependent manner and can assemble into several different AMPK heterotrimer complexes (2).

The AMPKγ-subunit plays an essential role for AMPK function by acting as the sensor of cellular adenosine nucleotide levels. The C-terminal end of AMPKγ is rather uniform between AMPKγ1-3 and includes four cystathionine β-synthase tandem repeats (CBS 1-4). These repeats form binding pockets for AMP, ADP and ATP (2,3). During increased ATP turnover e.g., during muscle contraction, AMP and ADP accumulate in the muscle cell facilitating their binding to AMPKγ. This activates AMPK allosterically, but also increases covalent activation of AMPK through initiation of conformational changes that alter the accessibility to AMPKα for upstream kinase(s) and phosphatase(s) (1,4).

In human skeletal muscle, three AMPK heterotrimers are found (AMPKα2β2γ1, α2β2γ3 and α1β2γ1)(5), whereas two additional (α2β1γ1 and α1β1γ1) are found in mouse skeletal muscle (6). Regardless of species, the AMPKγ3 protein is only expressed in glycolytic skeletal muscle (7–11). Despite this unique expression profile, the mechanistic importance of AMPKγ3 remains largely unexplored in human skeletal muscle.

A naturally occurring mutation of human AMPKγ3 arginine 225 to tryptophan (R225W) has previously been identified in members of two independent families (12). The R225W mutation is placed in the CBS 1 domain and was reported to elevate the total pool of AMPK activity in cultured cells from AMPKγ3 R225W carrier muscle. After the initial description of this mutation, it has been proposed that different AMPK heterotrimers exhibit various patterns of activity and activation patterns (5,13,14), so since the point mutation was found in AMPKγ3, one could speculate the change in total AMPK activity to originate from elevated basal activity of AMPKα2β2γ3 in AMPKγ3 R225W carrier muscle, but direct evidence supporting this remains unreported. Alternate mutations in the CBS-domains of AMPKγ-subunits have been described in human cardiomyocytes. Some are suggested to cause severe clinical disease, like Wolff-Parkinson-White syndrome, and the mutations are typically seen in combination with abnormal cardiac glycogen accumulation (15). Skeletal muscle from human AMPK R225W carriers was reported to contain twice the amount of glycogen as compared to controls (12). Thus, this proposes a link between AMPKγ-mutations and regulation of glycogen stores.

In the present study, we gained access to muscle biopsies from three AMPKγ3 R225W human carriers and individually matched controls enabling us to investigate how this mutation affects the heterotrimer-specific AMPK activity in mature skeletal muscle. To illuminate any causative links between the R225W mutation and further phenotypical characteristics, we generated a novel knock-in mouse model carrying the R225W mutation. We hypothesized that skeletal muscle carrying the human AMPKγ3 R225W mutation expresses high AMPKα2β2γ3 activity and that this would correlate positively with muscle glycogen levels.

## RESULTS

### AMPKα2β2γ3 activity and glycogen levels are not elevated in mature skeletal muscle of human AMPKγ3 R225Wcarriers

We obtained resting muscle biopsies from three individuals carrying the AMPKγ3 R225W mutation (R225W MUT) and three individually matched controls. Age, anthropometrics and clinical measures were obtained in controls and R225W MUT (Fig. S1). From the muscle biopsies, we isolated intact AMPK heterotrimers through sequential immunoprecipitations and measured kinase activity with or without exogenous AMP present in the assays. Within all pairs, the AMPKα2β2γ3 activity, in the absence of AMP, was lower in R225W MUT compared to controls. When AMP was present, the activity was similar between genotypes (Fig. 1A). The activity of AMPKα2β2γ1 and AMPKα1β2γ1 were comparable between groups, and the addition of AMP to the assay increased the activity of both heterotrimers independent of genotype (Fig. 1B-C). In contrast to our hypothesis, this suggests that the AMPKγ3 R225W mutation does not increase basal activity of the AMPKα2β2γ3 complex in mature human skeletal muscle.

**Fig. 1.**
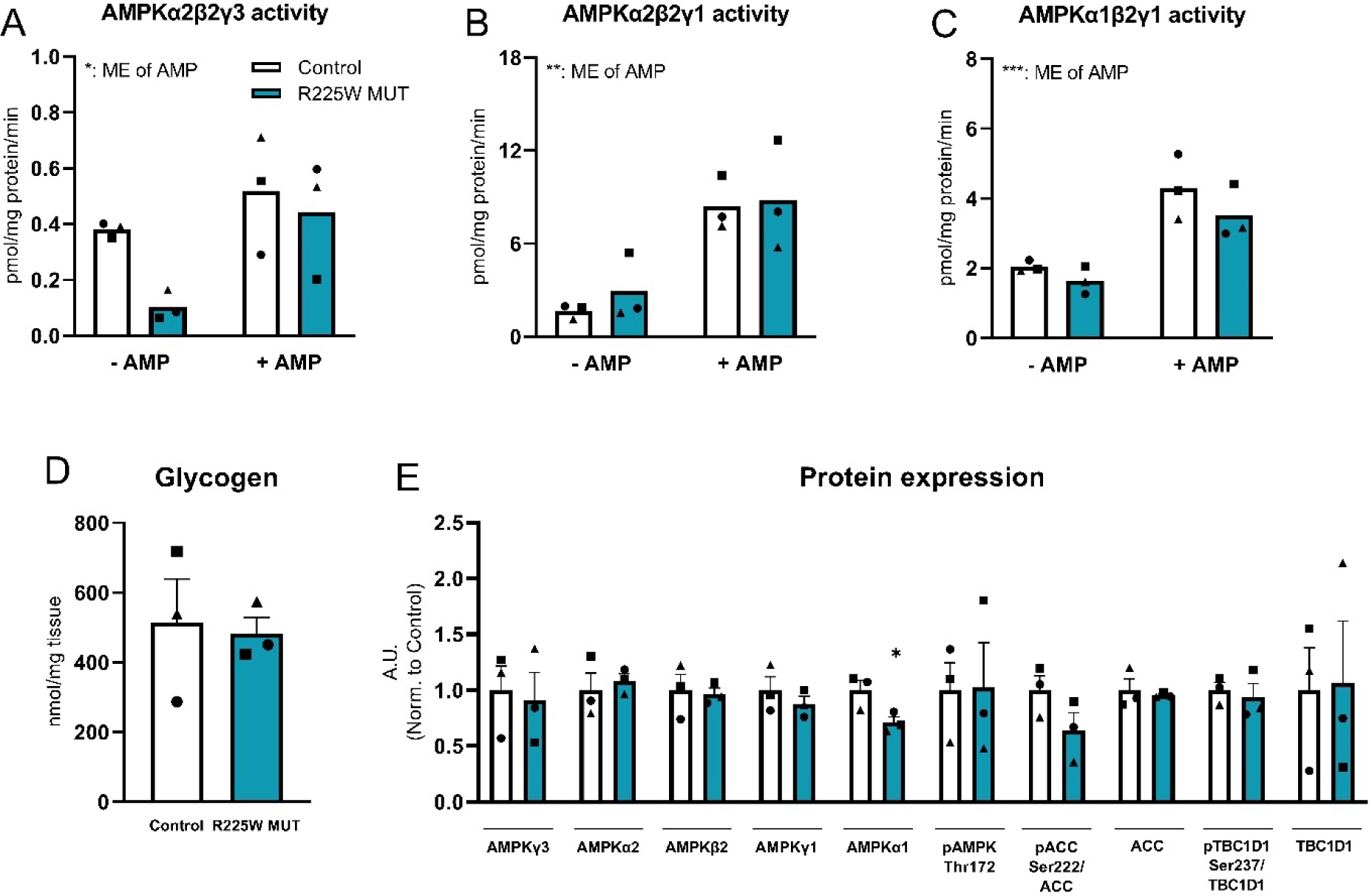
AMPK activity, AMPK expression and glycogen levels in skeletal muscle of human AMPKγ3 R225W carriers and controls. Muscle biopsies from *m. vastus lateralis* were obtained from human carriers of the naturally occurring AMPKγ3 R225W mutation (R225W MUT, teal bar) and individually matched controls (Control, white bar). Activity of AMPKα2β2γ3 (**A**), AMPKα2β2γ1 (**B**) and AMPKα1β2γ1 (**C**) were measured in muscle with or without addition of 200 µM AMP in the activity assay (n=3). Glycogen levels were measured in muscle (**D**, n=3). Protein expressions and phosphorylated levels of the indicated proteins and phosphorylation sites were measured in muscle and normalized to control level for each protein (**E**, n=3). Data are given as mean + SEM and individual values are included to identify the individually matched controls so that circle, square and triangle in the control group corresponds to the matching R225W MUT carrier. Two-way ANOVA (A-C) and unpaired t-test (D-E) were used where applicable to statistically evaluate differences between genotypes, AMP-presence or the interaction. *: Indicates main effect of AMP (A-C) or difference within genotype (D-E): *: p<0.05, **: p<0.01, ***: p<0.001. A.U.: Arbitrary Units.

Due to the proposed link between AMPKγ3 R225W and glycogen accumulation, the glycogen levels from the same muscle biopsies were measured. The means of both groups were within the normal human physiological range but did not differ (Fig. 1D).

The protein expression levels of AMPKγ3, AMPKα2, AMPKβ2 and AMPKγ1 were comparable between genotypes, while AMPKα1 was reduced in R225W MUT muscle (Fig. 1E). In agreement with the AMPK activity measurements, phosphorylation of AMPK Thr172 and downstream target TBC1D1 Ser237 did not differ between genotypes while a trend towards reduced phosphorylation of ACC Ser222 in R225W MUT muscle was apparent from the data, despite not reaching statistical significance (Fig. 1E). This suggests that the AMPKγ3 R225W mutation does not increase endogenous activity of the AMPKα2β2γ3 complex in mature human skeletal muscle, if anything, it associates with decreased activity.

### Homozygote AMPKγ3 R225W KI mice show suppressed AMPKα2β2γ3 activity and expression of AMPKγ3, while glycogen remains unchanged in skeletal muscle

To gain further understanding of the importance of the AMPKγ3 R225W mutation, we studied mice that were homozygous for this mutation. No significant difference was observed in body weight between adult wild-type (WT) and homozygous AMPKγ3 R225W knock-in (KI HOM) mice (Fig. 2A). KI HOM mice expressed a slightly leaner phenotype (Fig. S2A-B) despite eating more and oxidizing more carbohydrates in the dark phase (Fig. S2C-D). WT and KI HOM mice had similar activity patterns in the cage (Fig. 2B), during wheel running (Fig. S2G) and ran the same distance when running to exhaustion on treadmills (Fig. 2C, S2E-F). Overall, whole-body assessments did not unveil any substantial metabolic or behavioral phenotype of the KI HOM model.

**Fig. 2.**
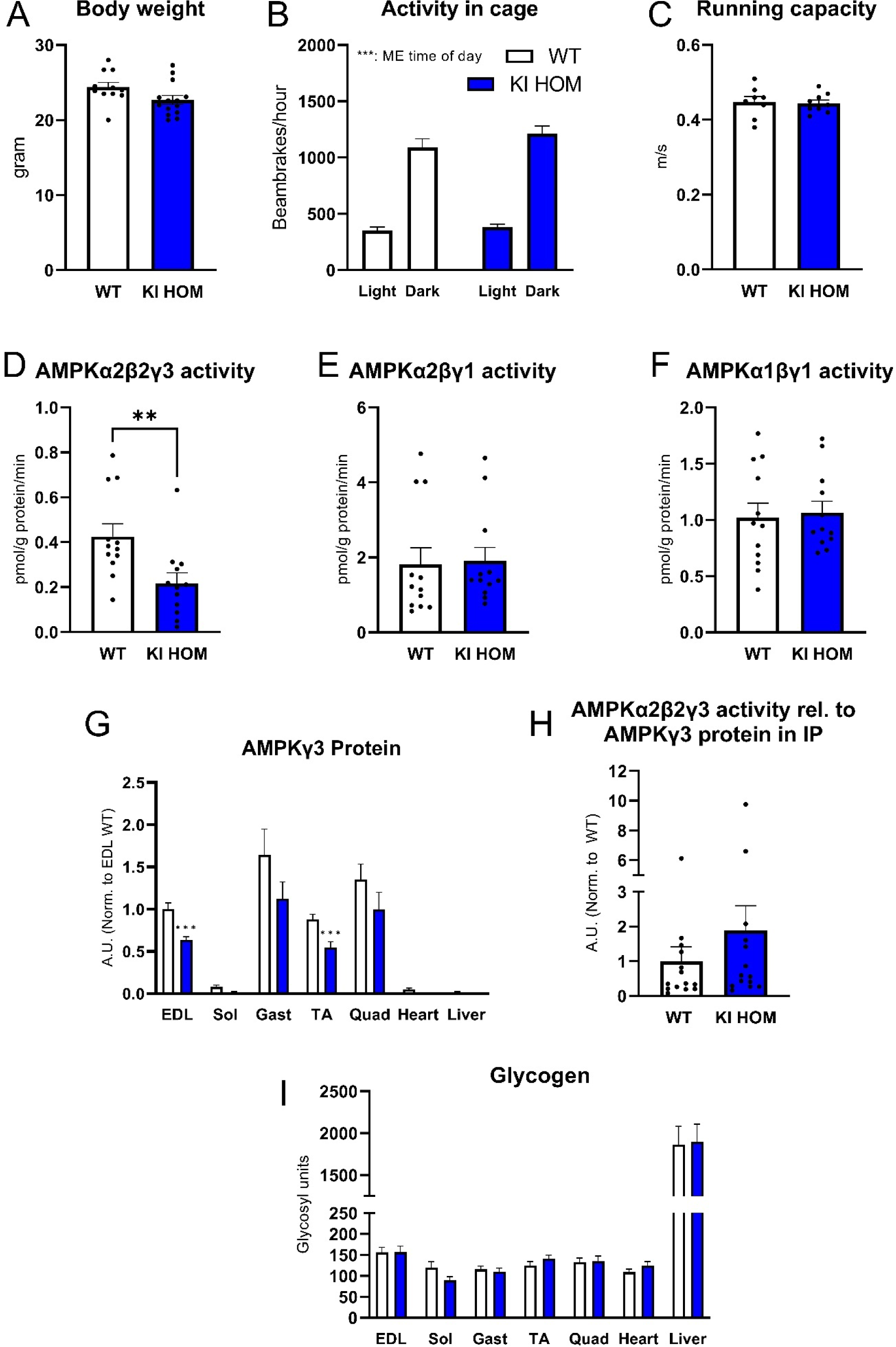
Whole-body characteristics, AMPK activity and glycogen levels in skeletal muscle of mouse homozygote AMPKγ3 R225W carriers and WT. Wild-type (WT, white bar) and homozygote AMPKγ3 R225W carrier (KI HOM, blue bar) mice were investigated on a whole-body and muscle level in the basal state. Body weight (**A,** n=11). Activity in cage (**B**, n=11-13). Maximal running capacity (**C,** n=8-9). Activity of AMPKα2β2γ3 **(D)**, AMPKα2βγ1 **(E)** and AMPKα1βγ1 **(F)** was measured in *m. tibialis anterior* with 200 µM AMP added to the activity assay (n=12). **G:** Protein expression of AMPKγ3 was measured in the indicated isolated muscles and tissues. Results are normalized to WT EDL levels for comparison (n=11). AMPKα2β2γ3 activity was measured in a separate set of *m. tibialis anterior* and related to the AMPKγ3 protein expression found in the immunoprecipitation (**H**). Data are normalized to WT level (n=14-15). Glycogen levels were measured in the indicated isolated muscles and tissues (**I**, n=6-18). Data are given as mean + SEM including dots to indicate individual values where applicable. Unpaired t-test (A, C-I) and Two-way ANOVA (B) were used where applicable to statistically evaluate differences between genotypes, time of day or the interaction. *: Indicates main effect of genotype/time of day: **: p<0.01, ***: p<0.001. A.U.: Arbitrary Units. EDL: m. e*xtensor digitorum longus*. Sol: *m. soleus*. Gast: *m. gastrocnemius*. TA: *m. tibialis anterior*. Quad: *m. quadriceps*. ME: Main Effect.

Measurements of AMPK heterotrimer-specific activity from isolated *m. tibialis anterior* (TA) displayed a 50% suppression of basal AMPKα2β2γ3 activity in KI HOM compared to WT controls (Fig. 2D), whereas no differences between genotypes with regards to AMPKα2βγ1 or AMPKα1βγ1 activities were observed (Fig. 2E-F). To understand this robust suppression of AMPKα2β2γ3 activity, the protein expression of AMPK subunits in several muscles and tissues from WT and KI HOM mice were measured. A lower expression of AMPKγ3 was evident in glycolytic muscles of KI HOM compared to WT. As expected, AMPKγ3 was not detected in *m. soleus* (Sol), heart or liver (Fig. 2G). Also, *PRKAG3* mRNA content did not differ in EDL and TA muscle from WT and KI HOM mice (Fig. S2N) suggesting post translational regulation of the AMPKγ3 protein expression level. There were no differences in protein expression of other AMPK subunits (Fig. S2H-L).

Since the method used for measuring heterotrimer-specific activity does not take AMPK subunit protein expression into account, we chose to correct for this in a separate set of samples, by relating the AMPKα2β2γ3 activity (Fig. S2O) to AMPKγ3 protein abundance in the immunoprecipitations (Fig. S2P). When doing so, we found no difference in muscle AMPKα2β2γ3 activity between WT and KI HOM mice. This indicates that the AMPKα2β2γ3 R225W heterotrimer is as active as the WT-form (Fig. 2H) but that the low abundance of AMPKγ3 R225W protein leaves the muscle cell *in vivo* exposed to less AMPKγ3-related activity. Glycogen levels measured in several different muscles, heart and liver tissue from resting WT and KI HOM mice were similar between genotypes (Fig. 2I).

### Insulin sensitivity of skeletal muscle is not affected in AMPKγ3 R225W KI HOM mice

Previous work from our group has indicated that increased AMPKγ3-related activity enhances insulin sensitivity of skeletal muscle (14,16). To evaluate the potential effects of AMPKγ3 R225W on insulin sensitivity, we subjected mice to a whole-body glucose-(GTT) and an insulin tolerance test (ITT) and investigated the response to insulin in isolated skeletal muscle *ex vivo*.

Fasting blood glucose concentrations were comparable between WT and KI HOM mice (Fig. 3A). The blood glucose concentration rose to a higher level in the KI HOM mice than in WT during the first 30 min of the GTT (Fig. 3A). The difference in blood glucose excursion faded after 60 min and the blood glucose remained concurrent over the last 60 minutes of the test (Fig. 3A). Plasma insulin levels did not change during the initial 30 minutes of the GTT in either genotype (Fig. 3B). Intraperitoneal insulin injection reduced the blood glucose concentration of both genotypes with no difference observed in the response throughout the test (Fig. 3C). In isolated and incubated *m. extensor digitorum longus* (EDL) and *m. soleus* (Sol), submaximal (100 µU/mL) and maximal (10.000 µU/mL) insulin-stimulation increased glucose uptake similarly between genotypes (Fig. 3D, G). The phosphorylation-state of central parts of the canonical insulin cascade i.e. pTBC1D4 Thr649 and pAkt Thr308 were comparable between genotypes for both EDL and Sol (Fig. 3E-F, H-I). This suggests that the whole-body glucose tolerance of KI HOM mice is slightly compromised, while the insulin sensitivity on a systemic and skeletal muscle level remain unaffected by the homozygous expression of AMPKγ3 R225W in skeletal muscle.

**Fig. 3.**
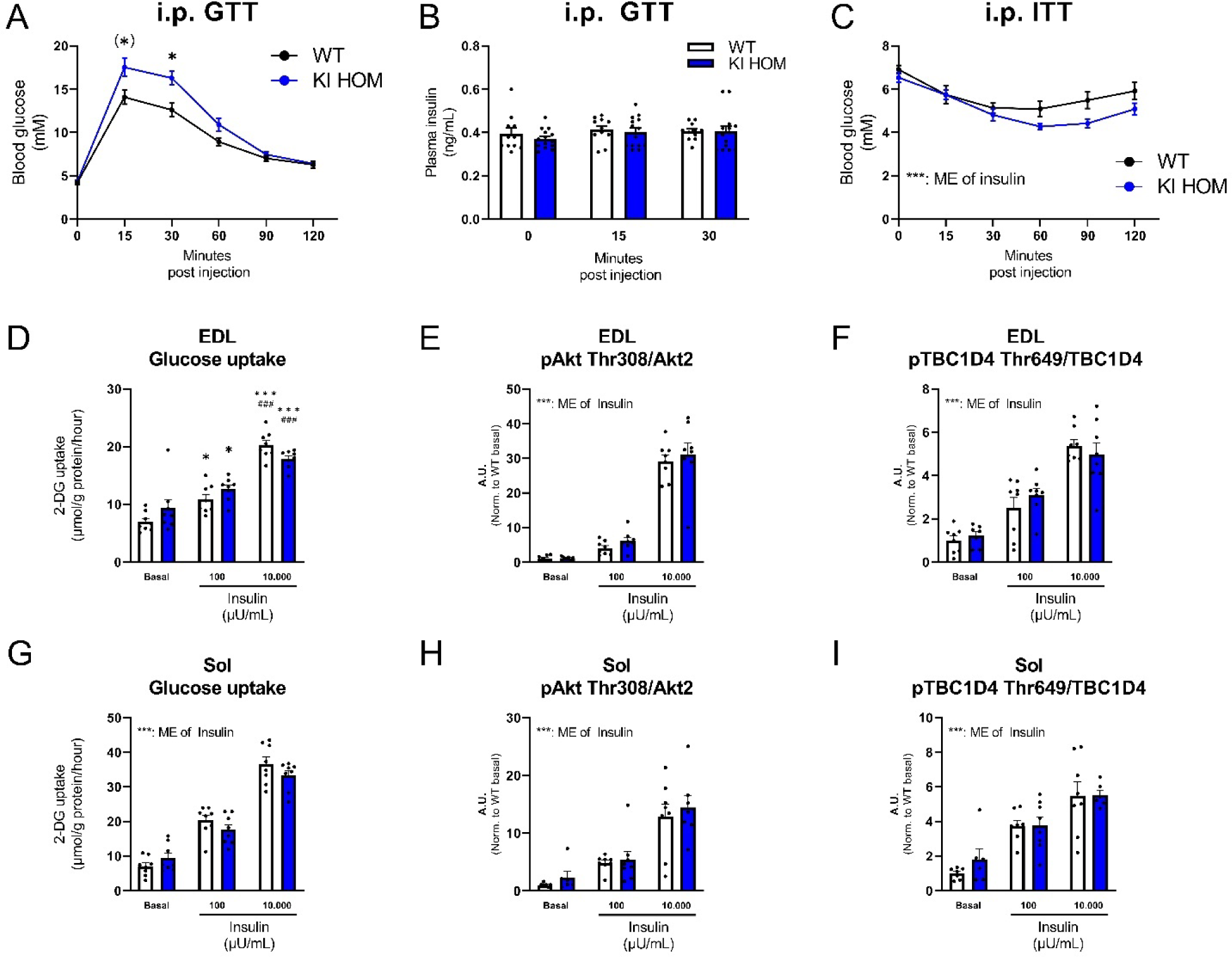
Whole-body and skeletal muscle insulin sensitivity of mouse homozygote AMPKγ3 R225W carriers and WT. The whole-body and muscle-specific insulin sensitivity of wild-type (WT, black line/white bar) and homozygote AMPKγ3 R225W carrier (KI HOM, blue line/bar) mice was investigated. Intraperitoneal (i.p.) glucose tolerance test (GTT) was performed. Blood glucose response from 0-120 minutes post glucose injection (**A**) and plasma insulin response (**B**) from 0-30 min post glucose injection in WT and KI HOM mice (n = 10-13). I.p. insulin tolerance test (ITT, 0.75 U/kg bodyweight) was performed in WT and KI mice (**C**). Blood glucose response from 0-120 minutes post insulin injection (n=10-13). Submaximal (100 µU/mL) and maximal (10.000 µU/mL) insulin stimulation was performed in isolated *m. extensor digitorum longus* (EDL; **D-F**) and *m. soleus* (Sol; **G-I**) excised from WT and KI HOM mice. Glucose uptake (D, G) and the phosphorylation level of pAkt Thr308 (E, H) and pTBC1D4 Thr649/TBC1D4 (F, I) was measured in EDL and Sol under basal or the two insulin-stimulated conditions *ex vivo*. All Western-blotting data are normalized to WT basal levels (n=8 both in EDL and Sol). Data are given as mean + SEM including dots to indicate individual values where applicable. Two-way ANOVAs with/without repeated measures and Holm-Sidak/Newman-Keuls post-hoc test (A, C-I) or unpaired t-tests (B) were used (where applicable) to investigate differences between genotypes, glucose/insulin or the interactions. *: (A-C) Indicates differences between genotypes at a given time point (D-I) Indicates differences compared to basal within genotype. #: indicates difference compared to submaximal insulin dose within genotype. *: p<0.05, ***/###: p<0.001. A.U.: Arbitrary Units. ME: Main Effect.

### The AMPKγ3 R225W mutation inhibits AICAR-induced AMPKα2β2γ3 activation and glucose uptake

A muscle incubation experiment using isolated EDL from WT and KI HOM mice was performed to evaluate the responses to the AMP-mimetic AICAR, which activates AMPK by binding to the γ-subunit CBS-domains. Both with and without AMP present in the *in vitro* activity assay, AMPKα2β2γ3 activity was increased with maximal AICAR-stimulation (2.0 mM) in WT EDL. This response was completely ablated in KI HOM muscle (Fig. 4A-B). In contrast, AICAR increased the activity of the AMPKα2βγ1 and AMPKα1βγ1 complexes in WT and KI HOM EDL, regardless of AMP presence (Fig. S4A-D). This indicates that AICAR-effects solely related to AMPKγ3 were affected by the AMPKγ3 R225W mutation.

**Fig. 4.**
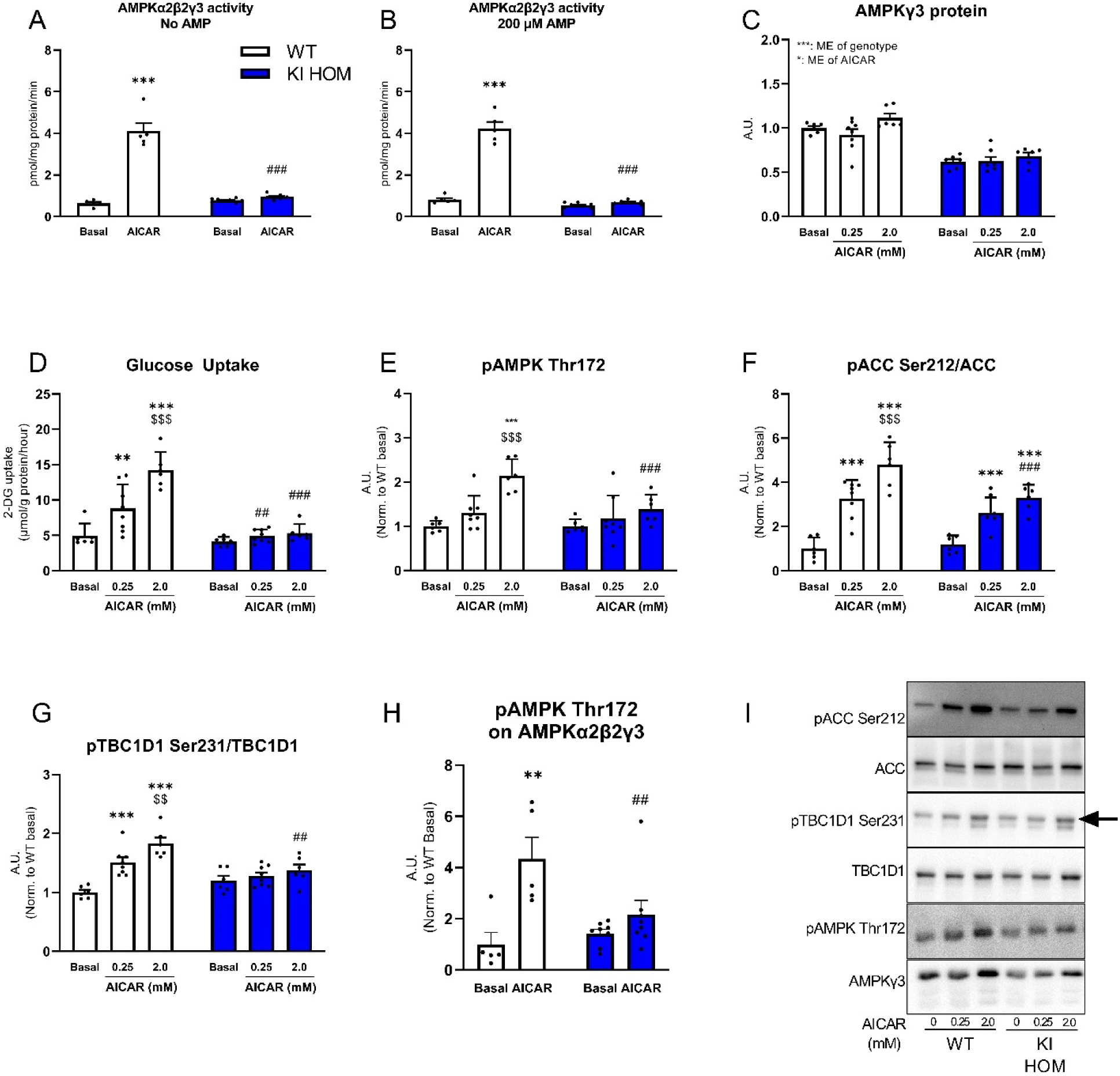
AMPK activity, glucose uptake and AMPK signaling in response to AICAR-stimulation in isolated muscle from mouse homozygote AMPKγ3 R225W carriers and WT. To understand the function of the AMPKγ3 R225W protein, we isolated *m. extensor digitorum longus* (EDL) from wild-type (WT, white bar) and homozygote AMPKγ3 R225W carrier (KI HOM, blue bar) mice and stimulated them with AICAR *ex vivo*. AMPKα2β2γ3 activity was measured in isolated EDL in the basal or AICAR-stimulated state (2.0 mM) without (**A**) or with (**B**) 200 µM AMP added to the activity assay (n=5-8). Protein expression of AMPKγ3 (**C**), glucose uptake (**D**) and phosphorylation levels of pAMPK Thr172 (**E**), pACC Ser212/ACC (**F**) and pTBC1D1 Ser231/TBC1D1 (**G**) were measured in incubated EDL in the basal state or under submaximal (0.25 mM) or maximal (2.0 mM) AICAR stimulation (n=6-8). The degree of pAMPK Thr172 specifically associated with AMPKα2β2γ3 was measured in AMPKγ3 immunoprecipitates from EDL stimulated with or without AICAR (**H**, **2**.0 mM) and related to the amount of AMPKα2 in the immunoprecipitates to correct for the different AMPKα2 protein levels. Representative blots relating to figure 4C-G (**I**). Data are given as means + SEM including dots to indicate individual values. All immunoblotting data is normalized to WT basal levels. Two-way ANOVA with Newman-Keuls post-hoc test was used (where applicable) to evaluate differences between genotype, AICAR or the interaction. *: Indicates effect of AICAR compared to basal within genotype. #: Indicates difference between genotypes within AICAR concentration. $: Indicates difference between AICAR concentrations within genotype. **/##/$$: p<0.01, ***/###/$$$: p<0.001. A.U.: Arbitrary Units. ME: Main Effect.

Prior studies by our group and others have found that AICAR-stimulated glucose uptake is dependent on AMPKγ3 in skeletal muscle (7,8,17). Therefore, we aimed to investigate the downstream impact of the R225W mutation on AMPKα2β2γ3 function by measuring glucose uptake during AICAR stimulation of isolated and incubated muscle. Glucose uptake was increased by submaximal (0.25 mM) and maximal (2.0 mM) AICAR stimulation in EDL muscle from WT mice, while the response to AICAR was absent in KI HOM EDL (Fig. 4D). AMPK Thr172 was phosphorylated in a dose-dependent manner in WT EDL whereas no effects of AICAR were observed in KI HOM EDL (Fig. 4E). Phosphorylation of The ACC Ser212 was increased in a dose-dependent manner in EDL from both WT and KI HOM mice, although the degree of phosphorylation was lower in KI HOM muscles compared to than WT muscles (Fig. 4F, S4E). The phosphorylation of the AMPK downstream target TBC1D1 Ser231 was dose-dependently increased by AICAR in WT muscle, but absent in KI HOM muscle (Fig. 4G, S4F).

To investigate if the absent phosphorylation of AMPK Thr172 on AMPKα is associated with the AMPKα2β2γ3 R225W complexes, phosphorylation of AMPKα2 Thr172 was measured in AMPKα2β2γ3 complexes immunoprecipitated from WT and KI HOM muscle and related to the amount of AMPKα2 protein in the IPs. AICAR increased the phosphorylation of AMPKα Thr172 in AMPKα2β2γ3 complexes from WT EDL, while the effect of AICAR was absent in KI HOM EDL (Fig. 4H, S4G-H). H). Taken together, the effects of AICAR on AMPKα2β2γ3 activation, glucose uptake and cellular signaling are impaired in isolated skeletal muscle from homozygote AMPKγ3 R225W KI mice.

### The AMPKγ3 R225W mutation inhibits contraction-induced activation of AMPKα2β2γ3 despite comparable glycogen utilization

We speculated if a more physiological and complex stimulus would be sufficient to activate the AMPKα2β2γ3 R225W heterotrimers. We have previously shown that AMPKα2β2γ3 activity is robustly increased in mouse muscle by *in situ* contractions (14). Applying this stimulus to WT and KI HOM mice, we found that the WT AMPKα2β2γ3 complexes were activated by muscle contractions. In contrast, the response was completely ablated by the AMPKγ3 R225W mutation both in the absence and presence of AMP (Fig. 5A-B). With no AMP in the *in vitro* assay, only AMPKα2βγ1 in WT muscle was activated by contractions (Fig. S5A), while contractions activated AMPKα2βγ1 in both genotypes when the assay included AMP (Fig. S5B). AMPKα1βγ1 was not activated by muscle contractions in either genotype (Fig. S5C-D). Despite the lack of AMPKα2β2γ3 activation in KI HOM muscle, contraction-induced glucose uptake was comparable between genotypes (Fig. 5D). This underscores that AMPK, and AMPKγ3 in particular, is not necessary for glucose uptake during contractions but rather acts to regulate muscle glucose uptake in recovery from contractions and exercise (17–19). In parallel, glycogen utilization during contractions was equal between genotypes (Fig. S5E). The regulation of AMPK Thr172 and ACC Ser212 was increased by contraction both in WT and KI HOM muscle, however, the increase was more pronounced in WT muscle (Fig. 5E-F, S5G). The phosphorylation of TBC1D1 Ser231 was increased to the same extent in the two genotypes (Fig. 5G, S5F). A similar contraction-induced phosphor-regulation of Erk1/2 Thr202/Tyr204 and p38 Thr180/Tyr182 indicates comparable induction of cellular metabolic stress during contraction in the two genotypes (Fig. S5H-K). To evaluate how much of the phosphorylation of AMPKα Thr172 was related to AMPKγ3, we immunoprecipitated AMPKγ3 and measured the contraction-induced phosphorylation of the co-immunoprecipitated AMPKα2. The Thr172 phosphorylation associated with AMPKα2 was increased vastly by contraction in the IPs from WT muscle but was completely absent in KI HOM IPs (Fig. 5H, S5L-M). Taken together, these findings suggest that the impaired activation of the AMPKα2β2γ3 complex is linked to less Thr172 phosphorylation of AMPKα2 and that enhanced activity associated with AMPKγ3 during contraction is not necessary for contraction-induced glucose uptake.

**Fig. 5.**
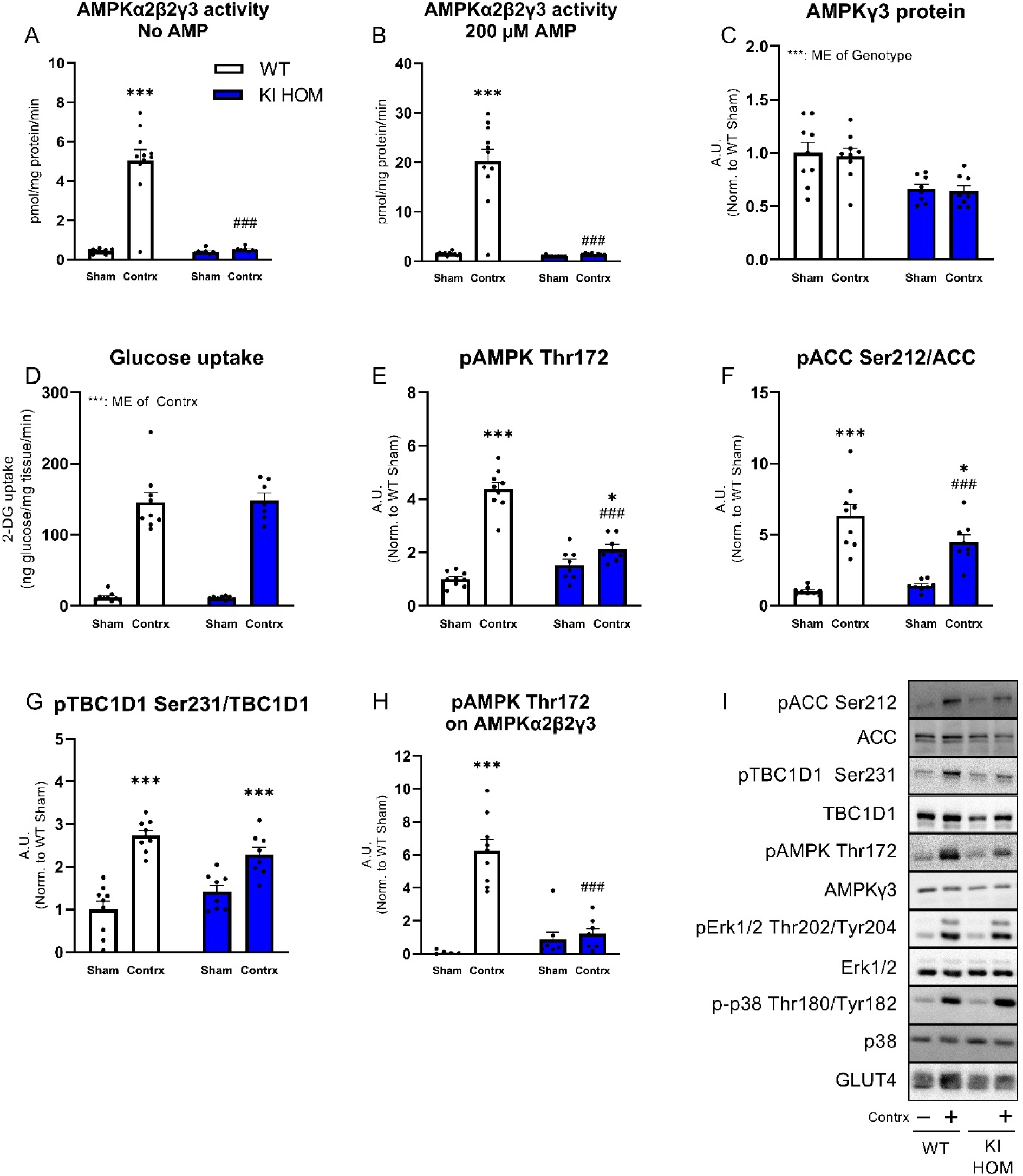
AMPK activity, glucose uptake and AMPK signaling in response to muscle contraction in muscle from mouse homozygote AMPKγ3 R225W carriers and WT. To understand the function of the AMPKγ3 R225W protein, we contracted one lower hindlimb and isolated *m. tibialis anterior* (TA) from wild-type (WT, white bar) and homozygote AMPKγ3 R225W carrier (KI HOM, blue bar) mice and left the contralateral leg as a sham-operated control. AMPKα2β2γ3 activity was measured in the sham-operated or contracted state without (**A**) or with (**B**) 200 µM AMP added to the activity assay (n=8-11). Protein expression of AMPKγ3 (**C**), glucose uptake (**D**) and phosphorylation levels of pAMPK Thr172 (**E**), pACC Ser212/ACC (**F**) and pTBC1D1 Ser231/TBC1D1 (**G**) were measured in TA (n=8-9). The degree of pAMPK Thr172 specifically associated with AMPKα2β2γ3 was measured in AMPKγ3 immunoprecipitates from TA in Sham or Contrx related to the amount of AMPKα2 in the immunoprecipitates to correct for the different AMPKα2 protein levels (**H**). Representative blots relating to figure 5C-G (**I**). Data are given as means + SEM including dots to indicate individual values. All immunoblotting data is normalized to WT Sham levels. A two-way ANOVA with Newman-Keuls post-hoc test was used (where applicable) to evaluate differences between genotype, contraction or the interaction. *: Indicates effect of contraction compared to sham within genotype. #: Indicates difference between genotypes within contraction-state. *: p<0.05, ***/###: p<0.001. A.U.: Arbitrary Units. ME: Main Effect.

### Heterozygote expression of AMPKγ3 R225W in mice mildly rescues effect of AMPKγ3

Human carriers and KI HOM mice are hetero- and homozygous for the AMPKγ3 R225W mutation, respectively. Based on previous findings (12) it is possible that the concurrent expression of AMPKγ3 WT and AMPKγ3 R225W protein in muscle from human carriers can influence the overall AMPKα2β2γ3 activity pattern. Therefore, we chose to study a mouse model heterozygous for the AMPKγ3 R225W mutation (KI HET) that expresses both AMPKγ3 WT and R225W protein.

In the KI HET mouse model, the expression of AMPKγ3 protein in EDL was only marginally suppressed compared to WT levels, while the protein, as expected, was absent in Sol (Fig 6A). When measuring AMPK heterotrimer-specific activity in isolated EDL muscle in the basal state, AMPKα2β2γ3 activity tended to be lower in the KI HET muscle compared to WT muscle (p=0.08, Fig. 6C), likely a reflection of the lower AMPKγ3 protein abundance. During maximal AICAR-stimulation (2.0 mM) AMPKα2β2γ3 was activated in both WT and KI HET EDL, but the AICAR effect in KI HET EDL was markedly blunted (Fig. 6C). Neither AMPKα2βγ1 nor AMPKα1βγ1 activity was increased by AICAR stimulation (Fig. S6A-B). Thus, the effect of heterozygote expression of AMPKγ3 R225W, with regards to AMPKα2β2γ3 activity, was comparable between mouse and human heterozygous carriers (Fig. 1A).

**Fig. 6.**
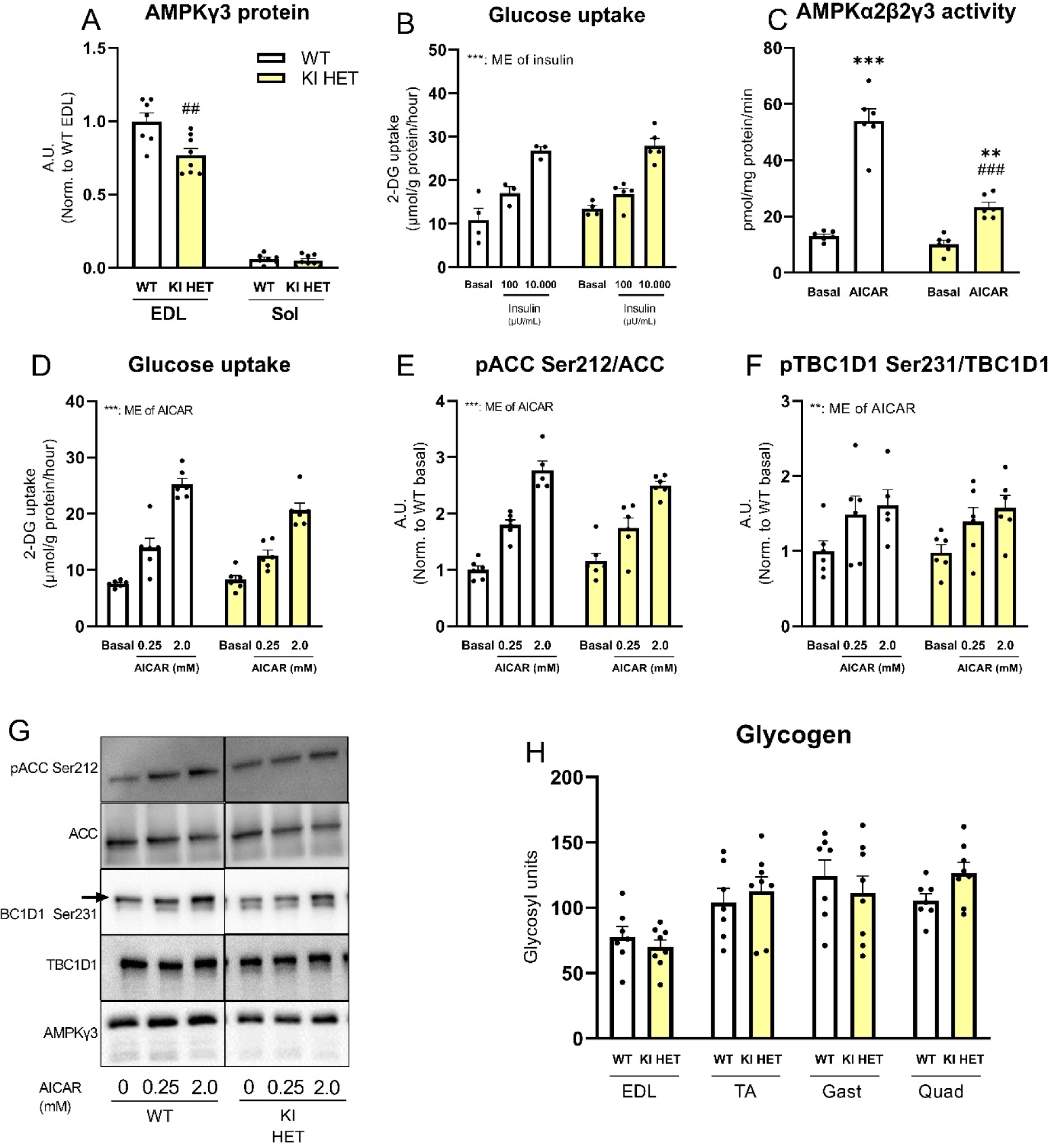
Insulin sensitivity, response to AICAR-stimulation and glycogen levels in isolated muscle from mouse heterozygote AMPKγ3 R225 carriers and WT. To introduce a combination of AMPKγ3 WT protein and AMPKγ3 R225W, we developed heterozygote AMPKγ3 R225W carrier mice (KI HET, yellow bar) and compared them to wild-type mice (WT, white bar). Protein expression of AMPKγ3 in *m. extensor digitorum longus* (EDL) and *m. soleus* (Sol) isolated from mice in basal condition (**A**, n=7-8). The glucose uptake of isolated EDL in the basal state or under submaximal (100 µU/mL) or maximal (10.000 µU/mL) insulin stimulation (**B**, n=3-5). AMPKα2β2γ3 activity was measured in isolated EDL in the basal or AICAR-stimulated state (2.0 mM) with 200 µM AMP added to the activity assay (**C**, n=6). Glucose uptake (**D**) and the phosphorylation levels of pACC Ser212/ACC (**E**) and pTBC1D1 Ser231 (**F**) were measured in incubated EDL in the basal state or under submaximal (0.25 mM) or maximal (2.0 mM) AICAR stimulation (n=5-6). Representative blots are given (**G**, the two genotypes were run on same gels/membranes). Glycogen levels were measured in the indicated isolated muscles (**H**, n=7-8). Data are given as mean + SEM including dots to indicate individual values. Unpaired t-test (A+H) or two-way ANOVA with or without Newman-Keuls post-hoc test (B-F) were used (where applicable) to investigate differences between genotype, insulin/AICAR or the interaction. *: Indicates effect of insulin/AICAR. #: Indicates effect of genotype. **/##: p<0.01, ***/###: p<0.001. A.U.: Arbitrary Units. ME: Main Effect.

The function of AMPKγ3 was assessed in KI HET by measuring *ex vivo* AICAR-stimulated glucose uptake. A comparable effect of AICAR on glucose uptake in EDL from WT and AMPKγ3 KI HET was observed (Fig. 6D). This pattern was also apparent in the downstream signaling of AMPK, where ACC Ser212 and TBC1D1 Ser231 were phosphorylated to the same extent between genotypes (Fig. 6E-F).

Importantly, the glycogen levels in muscle of KI HET mice corresponded to that of WT counterparts. (Fig. 6H).

Collectively, these findings suggest that the concurrent expression of WT AMPKγ3 and AMPKγ3 R225W does not potentiate any regulation in skeletal muscle beyond what could be expected from the partial introduction of WT AMPKγ3 protein.

## DISCUSSION

Here we report the heterotrimer-specific AMPK activity measured in mature skeletal muscle of human carriers of the AMPKγ3 R225W mutation. As AMPKγ-subunits are sensitive to AMP, we investigated the heterotrimer-specific activity both with and without AMP in our activity assay. In contrast to our hypothesis, we found lower AMPKα2β2γ3 activity in R225W MUT muscle in the absence of AMP, while AMP addition ‘normalized’ the activity to what was observed in muscle of control subjects. These findings do not resemble the initial reports of increased total AMPK activity in cultured muscle cells derived from the AMPKγ3 R225W carriers (12). The conflicting results could be due to methodological differences. Firstly, in the present study we measure the AMPK activity specifically associated with AMPKγ3 and not – as by Costford and colleagues – total activity of partially purified AMPK (12). Since AMPKα2β2γ3 corresponds to ∼20% of the AMPK pool in human skeletal muscle (5), this is a relevant distinction. From our prior studies, we know that AMPKα2β2γ3 activity only accounts for a marginal fraction of total AMPK activity, when the muscle is in the resting non-stimulated state. Accordingly, differences in basal AMPKα2β2γ3 activity are difficult to decipher from total AMPK activity measurements and would possibly be concealed by the basal activity of the other AMPK heterotrimer complexes. Secondly, for our measurements we use mature skeletal muscle and not – as by Costford and colleagues – differentiated myotubes. It is well known that expression of AMPK subunits differs markedly between myotubes and skeletal muscle, which means that the stoichiometry of AMPK subunits, and therefore the complex composition, varies substantially (20). As measurements comparing the protein expression of all AMPK subunits were not reported in the myotubes, this further complicates interpretation. Lastly, in the previous work, AMPK was only partially, and the activity assay included the SAMS-peptide as kinase substrate (12). As SAMS-peptide is based on a pACC1/2 Ser79/Ser212-consensus motif and ACC can be phosphorylated by other kinases (18,21), the data obtained using partially purified AMPK may not exclusively reflect AMPK activity. That being said, it is possible that subject heterogeneity within groups of the present cohort may vary from the previous cohort, which could account for some differences between studies. Furthermore, since we and the previous report, studied a rare mutation, the sizes of both our and the previous cohort were small, which certainly limits the strength of the findings. Therefore, we included studies of an AMPKγ3 R225W knock-in mouse model to strengthen our human investigations.

From these studies, we found that the AMPKγ3 R225W mutation does not affect the basal activity of AMPKα2β2γ3 with AMP present in the *in vitro* assay. This resembles what we demonstrated in muscle of the human carriers. Studies overexpressing AMPKγ3 R225Q in cell lines have shown both unchanged and increased effects on basal heterotrimer activity (17,22), while overexpression of AMPKγ3 R225Q in mouse skeletal muscle has been reported to reduce the AMPKγ3-associated activity, when comparing to muscle overexpressing AMPKγ3 WT protein. Of note, our initial measurements of AMPKα2β2γ3 activity were complicated by the apparent lower AMPKγ3 protein expression in KI HOM muscle. Based on prior analyses, we know that our immunoprecipitation procedure depletes the muscle lysate for AMPKγ3 protein in both WT and KI HOM mice. Therefore, we corrected for the altered protein expression by relating the activity to the AMPKγ3 protein in the immunoprecipitations. In previous reports, controls of this manner have not been carried out and the supraphysiological levels of AMPKγ3 R225Q protein may have influenced measurements of AMPK activity (11,17,23). Thus, we interpret this as the AMPKγ3 R225W-form does not affect the basal activity of the AMPKα2β2γ3 heterotrimer *per se*, but, at least in KI HOM, rather affects the expression of AMPKγ3 protein and consequently, the absolute amount of cellular AMPKα2β2γ3 activity. In muscle of the human carriers, heterozygous for AMPKγ3 R225W, we detect a comparable AMPKγ3 protein expression profile, manifesting in the similar basal activity with control subjects in the presence of AMP. Our data from the KI HET mice further supports this notion, as the introduction of AMPKγ3 WT protein rescued the basal activity to levels comparable to WT muscle. Thus, evidence from studies of mature resting muscle supports that AMPKγ3 R225W does not affect AMPKα2β2γ3 activity, and these *in vitro* measurements are reflected *in vivo* as downstream targets ACC and TBC1D1 are phosphorylated to the same extent during resting/non-stimulated conditions.

The mutation ablates activation of the AMPKα2β2γ3 heterotrimer by AICAR. As AICAR is metabolized to ZMP in muscle (24), our findings suggest that binding of ZMP to CBS 1 is compromised. AMP/ZMP binding to the CBS-domains works to potentiate AMPK activity in part by changing the access to AMPKα Thr172 for upstream kinases and phosphatases (4). This concept seems fully reflected in the observation that AICAR-induced AMPKα2 Thr172 phosphorylation is absent in KI HOM muscle. We, and others, have shown that AMPKγ3 protein is necessary for AICAR-stimulated glucose uptake (7,8,17). The lack of AICAR-stimulated increase in glucose uptake in the present study therefore further supports that the CBS 1 domain of AMPKγ3 cannot bind ZMP. Previously, the effect of AICAR on glucose uptake was reported to be partially suppressed in isolated EDL muscle overexpressing AMPKγ3 R225Q (11,17). In these models, the degree of remnant endogenous AMPKγ3 WT protein was not reported. The partial effects of AICAR observed in those studies could be due to ZMP binding to the endogenous and ‘efficacious’ AMPKγ3 WT protein. In support, the effect of AICAR on glucose uptake was rescued in our study of KI HET muscle that partially expresses WT endogenous AMPKγ3 protein opposed to none in KI HOM. Thus, the sum of these data indicates that the AMPKγ3 R225W is malfunctioning. We propose that this relates, at least partially, to loss of nucleotide binding.

AMPKα2β2γ3 R225W complexes also remains inactive despite substantial glycogen utilization and stress-signaling during muscle contraction. Our findings indicate that the downstream canonical targets of AMPK are indeed phosphorylated, though to a lower degree in KI HOM muscle, suggesting that only downstream regulation related to AMPKγ3-containing complexes is affected by the R225W mutation. In EDL overexpressing AMPKγ3 R225Q, overall AMPK activity was increased by *ex vivo* contraction albeit to a lesser extent than in WT muscle (23). As mentioned above, it is hard to decipher which AMPK heterotrimers (endogenous WT or R225Q overexpressed) are activated from overall AMPK measurements, but the study supports that mutations at the R225-position affects activation of the complete pool of AMPKγ3 heterotrimers (23). Taken together, these experiments propose that AMP-binding to CBS 1 is important for the full phosphorylation and activation of AMPKα2β2γ3 also during more complex physiological interventions. From structural and mutational studies *in vitro*, it has previously been suggested that AMP binding at the CBS 3 domain is of key importance for AMPKα Thr172 phosphorylation, while binding at the CBS 1 and CBS 4 domains is crucial for the affinity of CBS 3 for AMP (3). Thus, one model to explain the loss of nucleotide/AICAR-induced activation of AMPKα2β2γ3 could be that a mutation-induced impairment of AMP-binding in CBS 1 perturbs the important CBS 3 binding of AMP by reducing the affinity. Clearly, further studies are needed to investigate this directly. What this means for the human carriers of the R225W mutation remains an open question. At first glance, without performing statistical analyses, our data indicate that the AMP-sensitivity of human AMPKγ3 R225W is increased – contradicting what is found in the mouse work. However, it is important to bear in mind that the human carriers are heterozygous for the mutation and that the AMPKγ3 WT protein, found in muscle of the carriers, will likely respond to AMP accumulation.

Muscle glycogen accumulation is a hallmark finding in previous studies of human and animal mutations in AMPKγ3 R225 (11,12,17,23,25,26). Evident from our human data, we are not able to reproduce previous reports in human carriers of the AMPKγ3 R225W mutation (12) and the mouse models do not resemble what has been found in virtually all studies of the AMPKγ3 R200Q (naturally occurring in pigs, equivalent to mouse R225Q) or R225Q mutations (11,17,23,25–28). A clear difference between studies is the type of amino acid replacing arginine at position 225. No direct comparison of the R225Q and R225W has been made, but given that the properties of glutamine and tryptophan are quite different, e.g. with regards to polarity, their interaction with other amino acids or substances would likely vary. In relation to the human carriers, we cannot exclude that differences in methodology could influence our conflicting results. Nevertheless, it is striking that none of our mouse models express high glycogen levels in skeletal muscle, not even when the mutation is expressed homozygously.

The molecular targets of AMPKα2β2γ3 remain largely unknown. A study from our group has suggested that AMPKγ3 is involved in the regulation of muscle insulin sensitivity (16) and hyperinsulinemic euglycemic clamp experiments in two subjects indicate that the human carriers of the R225W mutation express higher whole-body insulin sensitivity (12). In line with the AMPKγ3 R225W mutation not causing activation of the heterotrimer, we did not observe any differences in insulin sensitivity for glucose uptake or central parts of the insulin-signaling cascade. Previous studies investigating insulin-stimulated glucose uptake in fed mice overexpressing the R225Q mutation did not find changes in insulin sensitivity either, further supporting our findings (11,17). This further goes hand in hand with the lack of glycogen phenotype seen in the models studied here.

## CONCLUSION

The AMPKγ3 R225W mutation does not impact AMPK-associated activity in mature human skeletal muscle and the mutation is not linked to glycogen accumulation, findings which are also reflected in the mouse models used here. Furthermore, the R225W mutation ablates AMPKγ3-associated activation by AICAR/muscle contractions, presumably through loss of nucleotide binding.

## MATERIALS AND METHODS

### Human subjects

The protocol for the human biopsies obtained from the *m. vastus lateralis* of human AMPKγ3 R225W carriers and matched controls was approved by the Research Ethics Board of the Ottawa Hospital and the University of Ottawa Heart Institute. Written consent was obtained from all participants. The control subjects were chosen to match each carrier on an individual level. Anthropometric and relevant clinical measures are given in figure S1.

### Animals

All animal experiments were carried out under approval from the Danish Animal Experiments Inspectorate (License #2019-15-0201-01659) and complied with the European Union guidelines for the protection of vertebrate animals used for scientific purposes. All mice were fed standard rodent chow and drinking water *ad libitum*. The mice were group housed on a 12:12 hour light-dark cycle in a temperature-controlled room (22 +/-2°C) in cages with nesting material unless otherwise stated. All mice used were adult females with an age spanning 10-34 weeks across experiments. Within experiments mice were age-matched between genotypes.

### Generation of AMPKγ3 R225W knock-in mice

AMPKγ3 R225W knock-in (KI) mice were generated at Jackson Laboratories (Bar Harbor, ME, USA) in the C57BL/6J background strain using the Prkag3exon6 sgRNA (CATGGTGGCCAACGGTGTGA) with the PRKAG3 R225W donor sequence (TCGTTCTTTCCTGCCCCTCAGATAAAGAAGGCTTTCTTTGCCATGGTGGCCAACGGTGTGTGGGCAGCTCCTCTGTGGGACAGCAAGAAGCAGAGCTTTGTGGGTGAGGAGAGGTGGCTGG). Founder animals were screened by PCR amplification and sequencing to confirm an insertion of the AMPKγ3 R225W mutation. Founder animals were bred with C57BL/6J mice for >8 generations prior to reported experiments. From the homozygote AMPKγ3 R225W KI mice, we also developed a mouse heterozygote for the AMPKγ3 R225W mutation, by crossbreeding homozygotes and wild-type mice. The heterozygosity was confirmed by PCR-based genotyping from earpieces.

### Body composition

Measurements of mice body compositions were obtained by magnetic resonance imaging scanning (EchoMRI 4-in-1TM, Echo Medical System LLC, Houston, TX, USA). Whole-body fat and lean mass were determined and related to total body weight.

### Basal calorimetry

Three days prior to testing, mice were single-housed in cages used for indirect gas calorimetry measurements (TSE, PhenoMaster metabolic cage systems, Bad Homburg, Germany). The mice had access to standard rodent chow and drinking water. Upon acclimatization, food intake, VO_2_-consumption, VCO_2_-production and habitual activity (measured as beam brakes) were recorded over 4 days. The respiratory exchange ratio (RER) was calculated from the VO_2_-consumption and CO_2_-production measurements (VCO_2_/VO_2_).

### Voluntary wheel-running

Mice were single-housed with access to running wheels, water and chow. Before measurements, the mice were allowed to acclimatize to the cage and running wheel for at least 2 days. Both individual running distance and running time were recorded over a 16-day period.

### Maximal running capacity

Mice were accustomed to treadmills over a 5-day period ranging from being placed on the treadmills for 10 minutes on day 1 to running for 12 minutes at 0.33 m/s at 0° incline on day 5. One week after treadmill adaptation, a maximal running capacity test was performed. The mice were single-housed, with access to chow and water *ad-libitum*, 3 days prior to the test. The test was performed by placing the mice on the treadmill, which gradually increased in speed from 0 to 0.16 m/s over 5 minutes at 15° incline. From here, the speed was increased with 0.02 m/s every minute maintaining constant incline. The mice ran until exhaustion, which was determined by a technician blinded for mouse genotype. Before and immediately after running cessation, venous blood glucose and lactate levels were measured from the tail.

### Glucose- and insulin tolerance tests

Mice were weighed and placed in individual cages with free access to drinking water. Prior to an intraperitoneal (i.p.) glucose tolerance test (GTT), mice were fasted overnight (16 hours). The mice were injected with 2 g/kg bodyweight of D-glucose dissolved in saline (20% glucose solution). Blood glucose was followed by venous blood samples taken from the tail 0, 15, 30, 60, 90 and 120 minutes after glucose injection. At the first 3 time points, blood samples were taken for subsequent determination of plasma insulin concentration by a standard insulin ELISA kit (ALPCO, NH, USA).

Prior to an i.p. insulin tolerance test (ITT), mice were fasted in individual cages for 4 hours. The mice were injected with 0.75 U/kg bodyweight of insulin (Actrapid, Novo Nordisk, Denmark) dissolved in physiological saline (0.9%). Blood glucose concentration was measured from the tail 0, 15, 30, 60, 90 and 120 minutes after insulin injection.

During both tests, blood glucose concentrations were determined using a glucometer (Contour XT, Bayer, Germany).

### Muscle incubation studies

Insulin- and AICAR-stimulated glucose uptake were determined in isolated skeletal muscle as previously described (29). In short, fed mice were anesthetized by injecting a mix of pentobarbital (100 mg/kg bodyweight), xylocaine (5.0 mg/kg bodyweight) and isotonic saline. *M. extensor digitorum longus* (EDL) and/or *m. soleus* (Sol) were dissected and suspended with suture loops on hooks in incubation chambers, at resting tension (Danish MyoTechnology, Denmark). The muscles recovered from dissection for at least 10 minutes in Krebs-Ringer buffer (KRB; 117 mM NaCl, 4.7 mM KCl, 2.5 mM CaCl_2_, 1.2 mM KH_2_PO_4_, 1.2 mM MgSO_4_, 24.6 mM NaHCO_3_, pH=7.4) supplemented with 0.1% bovine serum albumin, 2 mM Na-pyruvate and 8 mM mannitol, before they were stimulated with either basal, 100 µU/mL or 10.000 µU/mL insulin (Actrapid, Novo Nordisk, Denmark) for 30 minutes. Throughout the protocol, a gas mixture of O_2_ (95%) and CO_2_ (5%) was continuously supplied to the incubation chambers and the temperature of buffers were kept at 30°C. Furthermore, in a separate study, EDL muscles were isolated and treated with either basal, 0.25 mM or 2.0 mM AICAR (Toronto Research Chemicals, Canada) for 40 minutes. To determine glucose uptake, 1 mM [^3^H]2-deoxyglucose (0.028 MBq/mL) and 7 mM [^14^C]mannitol (0.0083 MBq/mL) were added to the incubation medium during the last 10 minutes of either insulin or AICAR stimulation. After incubation, the muscles were washed in cold (4°C) KRB, blotted dry and frozen in liquid nitrogen. Glucose uptake was determined from measurements performed on muscle lysate as previously described (29).

### In situ muscle contraction studies

Fed mice were anesthetized by injecting a mix of pentobarbital (90 mg/kg body weight), xylocaine (4.9 mg/kg bodyweight) and physiological saline. The fur on both hind limbs was removed, and the *n. peronei* were surgically exposed. An electrode was connected to the nerve of one leg, leaving the contralateral leg as rested sham-operated control. The muscles of the front side of the hind limb were contracted by applying 0.5 s trains (100 Hz, 0.1 ms, 5 V) every 1.5 second for a total of 10 minutes. To determine glucose uptake during contractions, a bolus of [^3^H]2-deoxyglucose (12.3 MBq/kg bodyweight) dissolved in physiological saline was injected retro-orbitally just prior to initiation of the contraction protocol. Venous blood samples were taken from the tail after 0, 5 and 10 minutes of muscle contraction. Skeletal muscle glucose uptake was assessed based on the amount of accumulated [^3^H]2-deoxyglucose-6-phosphate in *m. tibialis anterior* (TA) and tracer-specific activity in plasma from the blood samples as previously described (30).

### Tissue homogenization

Isolated mouse tissues (10-50 mg wet weight) were homogenized in 400 µL of cold (4°C) homogenization-buffer (10% glycerol, 50 mM HEPES, 1% NP40, 20 mM Na_4_P_2_O_7_, 150 mM NaCl, 2 mM PMSF, 1 mM EDTA, 1 mM EGTA, 3 mM benzamidine, 10 µg/mL leupeptin, 10 µg/mL aprotinin, 2 mM Na_3_VO_4_, pH=7.5) in Eppendorf tubes alongside cold steel beads. The tubes were shaken for 2×45 seconds at 30 Hz using a Tissue Lyser (TissueLyser II, Qiagen, Germany). After homogenization, 600 µL of homogenization-buffer was added to tubes containing either *m. gastrocnemius*, TA*, m. quadriceps*, heart or liver tissue. Next, the tubes were rotated end-over-end for one hour. For some analyses, tissue homogenates were used. This was collected after the rotation step. For other analyses using tissue lysates, the rotated tubes were centrifuged for 20 minutes at 16.000 g at 4°C. The homogenates/lysates were collected and frozen at −80°C until further analyses.

Human muscle biopsies were freeze-dried and dissected free of blood and fat tissue under microscope. The tissue was homogenized using the same buffer and procedure as above, but in sample-specific volumes of homogenization buffer (i.e., 1 mg dry weight to 250 µL buffer) to standardize protein concentrations in the resulting tissue lysate.

### Protein determination

The protein concentration of tissue homogenates and lysates was determined using the standard bicinchoninic acid method as described by the manufacturer (Thermo Fisher Scientific, MA, USA).

### SDS-PAGE and Western blotting

To investigate protein expression and molecular signaling in various tissues, we made use of SDS-PAGE and Western blotting. Proteins were separated by loading Laemmli-samples from tissue lysates in self-cast acrylamide gels (5-10%) and carried out SDS-PAGE. Prior to blotting, polyvinylidine difluoride (PVDF) membranes were activated in 96% ethanol for 5 minutes. Proteins were then transferred to the membranes by semi-dry Western blotting in a Trans-Blot Turbo Transfer System (Bio-Rad Laboratories Inc., CA, USA). Next, the membranes were blocked in TBST (10 mM Tris, 150 mM NaCl and 0.05% Tween20, pH=7.4) supplemented with either skimmed milk powder (2%) or bovine serum albumin (3%) for 5 minutes. The membranes were incubated in primary antibody on a rocking table at 4°C overnight. On the following day, the membranes were washed in TBST several times before incubation in secondary antibody for 45 minutes on a rocking table at room temperature. After additional washing in TBST, proteins were visualized using chemiluminescence and a digital imaging system (ChemiDoc XRS+; Bio-Rad, CA, USA).

### RNA isolation and reverse transcription

Total RNA was isolated from EDL and TA by a modified guanidinium thiocyanate-phenol-chloroform extraction method adapted from previous reports (31) as described previously (32) except that the tissue was homogenized for 120 seconds at 30 Hz in a Tissue Lyser (TissueLyser II, Qiagen, CA, USA). Superscript II RNase H^-^ system and Oligo dT (Invitrogen, CA, USA) were used to reverse transcribe the mRNA to cDNA, as described previously (32).

### Real-time PCR

The mRNA content of AMPKγ3 and TATA-box binding protein (TBP) was determined by real-time PCR using the QuantStudio 7 Flex detection system (Applied Biosystems, CA, USA) as previously described (33). TBP mRNA was determined by using a predeveloped assay reagent while primers and TaqMan probe were designed for detecting AMPKγ3 mRNA using the database www.ensembl.org and Primer Express 3.0 software (Applied Biosystems, CA, USA). Both real-time PCR reactions were performed using Universal MasterMix and TaqMan MGB probes with 5’FAM and a 3’ nonfluorescent quencher (NFQ, Thermo Fisher, CA. USA). The obtained cycle threshold values were converted to an arbitrary amount by using standard curves from a serial dilution of a pooled sample made from all samples. AMPKγ3 mRNA content was normalized to TBP mRNA content, which was not different between genotypes.

### Glycogen measurements

Due to scarceness of the human biopsy material, we measured glycogen content by dot blotting. Six µg lysate protein from each biopsy was spotted directly on a PVDF membrane and incubated with an antibody recognizing glycogen (34–36). The dots were visualized using secondary antibody and chemiluminescence and values were related to a standard curve made of a human muscle sample pool with a known glycogen content (measured biochemically).

Mouse tissue homogenate (500 µg) was boiled in 2M HCl for 2 hours in order for glycogen hydrolysis to occur. The resultant glycosyl units were measured using a fluorometric method (37).

### Heterotrimer-specific AMPK activity assay

Skeletal muscle heterotrimer-specific AMPK activity was measured as previously described (38). In short, 150 µg of lysate protein underwent sequential immunoprecipitations, aiming at precipitating first AMPKγ3, then AMPKα2 and lastly AMPKα1 from the same lysate sample, thus clearing the lysate from all AMPK complexes. In this way, the assay measures first the activity of AMPKα2β2γ3, then AMPKα2βγ1 and lastly AMPKα1βγ1. The activity assays were performed for 30 minutes including the immunoprecipitated complexes, AMARA-peptide and [^33^P_γ_]-labelled ATP tracer (Hartmann Analytic GmbH, Germany). The reactions were spotted on P81 filter paper (Saint Vincent’s Institute, Medical Research, Australia) and excess phosphate tracer was washed off the paper. The amount of [^33^P_γ_]-tracer on the paper (signifying the kinase activity) was analyzed using a Typhoon FLA 700 IP PhosphorImager (GE Healthcare, Denmark), and related to the specific activity of the kinase reaction buffer, which was measured using liquid scintillation counting. In an additional approach, we investigated the AMPKα2β2γ3 activity relative to the amount of AMPKγ3 protein within the genotype, by performing immunoprecipitations on 300 µg of protein from TA muscle and used one part (150 µg) for the AMPK activity assay and the other (150 µg) for determination of the AMPKγ3 protein amount through SDS-PAGE and Western-blotting (as described above).

### Antibodies used for Western blotting and immunoprecipitation

Primary antibodies against pACC Ser212/222 (3661, mouse/human), pAMPK Thr172 (2531), Erk1/2 (9102), pErk1/2 Thr202/Tyr204 (9101), p38 (9212), p-p38 Thr180/Tyr182 (9211), Akt2 (3063), pAkt Thr308 (9275) and pTBC1D4 Thr649 (8881) were from Cell Signaling Technology (Danvars, MA, USA). The primary antibody against pTBC1D1 Ser231/237 (07-2268, mouse/human) was from Millipore (Burlington, MA, USA) and the antibodies against TBC1D1 (ab229504), TBC1D4 (ab189890) and AMPKγ1 (ab32508) were from Abcam (Cambridge, UK). Antibodies against AMPKα2 (19131) and mouse AMPKβ1 (100357) were from Santa Cruz Biotechnologies (Dallas, TX, USA) while the antibody against GLUT4 was from Thermo Fisher Scientific (Waltham, MA, USA). Antibodies against AMPKα1 (Gift from Prof. Göransson, Lund University, Sweden), human AMPKβ2 (Gift from Prof. Hardie, University of Dundee, Scotland), human AMPKγ3 (GenScript, NJ, USA) and mouse AMPKγ3 (Yenzym, CA, USA) were custom made. ACC protein was detected using horseradish perodixase-conjungated streptavidin (Jackson ImmunoResearch, Cambridgeshire, UK).

The antibodies used for immunoprecipitations were all custom made; AMPKα1 at GenScript (NJ, USA), human AMPKα2 and AMPKγ3 at University of Dundee (MRC PPU Reagents and Services, Scotland) and mouse AMPKα2 and AMPKγ3 at Yenzym (CA, USA).

## Acknowledgements

The authors would like to thank Prof. Mary-Ellen Harper (University of Ottawa, ON, Canada) for providing the muscle biopsies of the human carriers and control subjects. The authors acknowledge the technical assistance from Irene B. Nielsen, Betina Bolmgren, Saif A. H. Al-Haidar and Ann-Sofie A. Kleis-Olsen (Department of Nutrition, Exercise, and Sports, Faculty of Science, University of Copenhagen, Denmark). Furthermore, we want to thank Russell Miller (Pfizer Inc., MA, USA) for providing the founder AMPKγ3 R225W KI mice and impactful scientific discussion.

## Author contribution

NOE, RK, CKP. and JFPW conceived and designed the research. NOE, RK, NSH, NRA, JBB, SR and HP performed the experiments. NOE analyzed the data and drafted the manuscript. RK and JFPW contributed to manuscript drafting. CKP provided the founder AMPKγ3 R225W KI mice. Every author contributed to data interpretation and critical discussion thereof as well as critical revision of the manuscript. All authors read and approved the final version of the manuscript. JFPW is the guarantor of the work and has full access to all data and takes responsibility for the integrity of the data and analyses provided.

## SUPPLEMENTARY MATERIALS

**Fig. S1.**
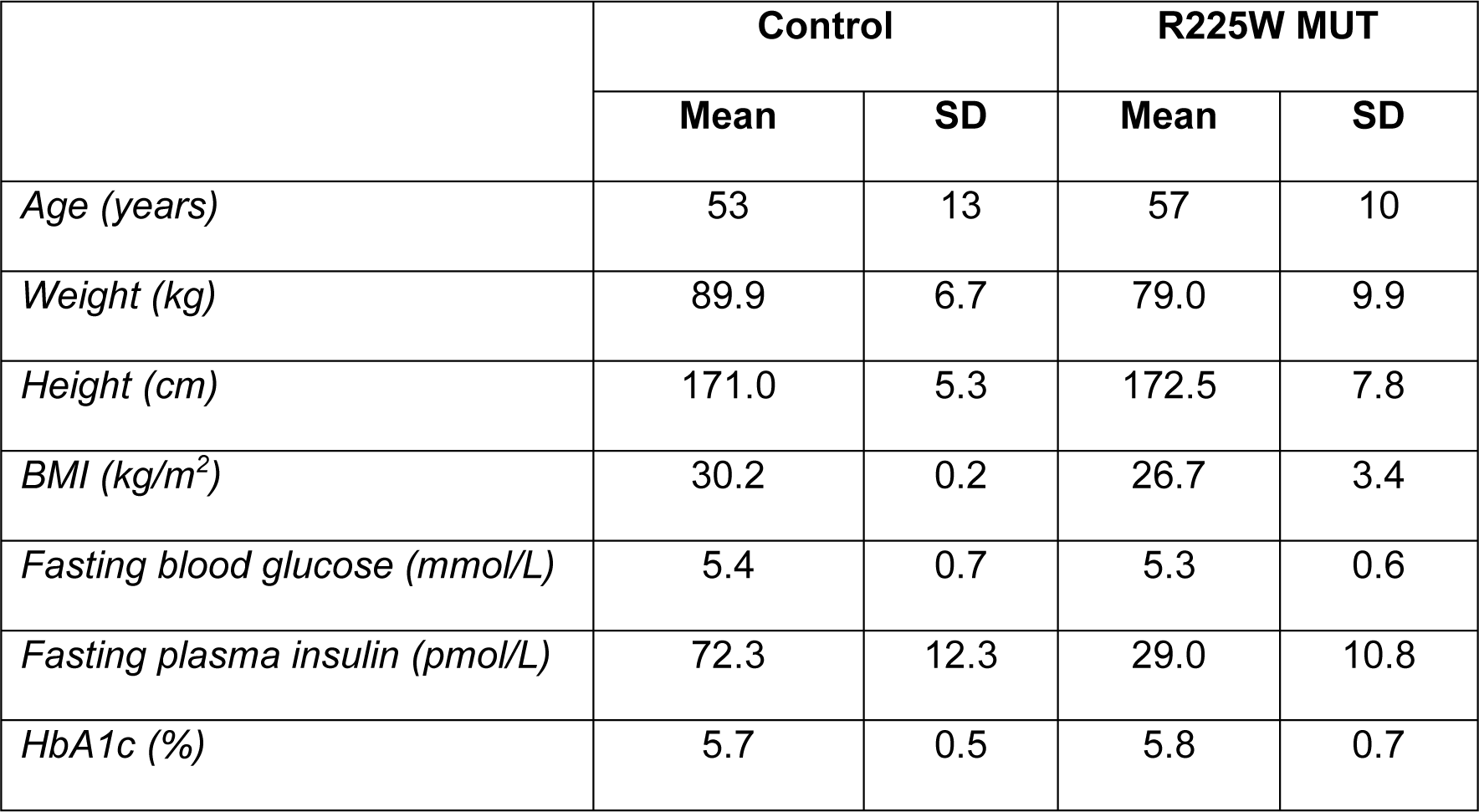
Age, anthropometrics and blood markers measured in human control and carriers of the AMPKγ3 R225W mutation. Age, anthropometrics and blood markers measured in human carriers of the AMPKγ3 R225W mutation (R225W MUT) and individually matched controls. Data are given as mean ± standard deviation (SD). Paried t-tests were performed to investigate differences between groups. No differences were observed. BMI: Body Mass Index, HbA1c: Glycated hemoglobin related to amount of total hemoglobin molecules.

**Fig. S2.**
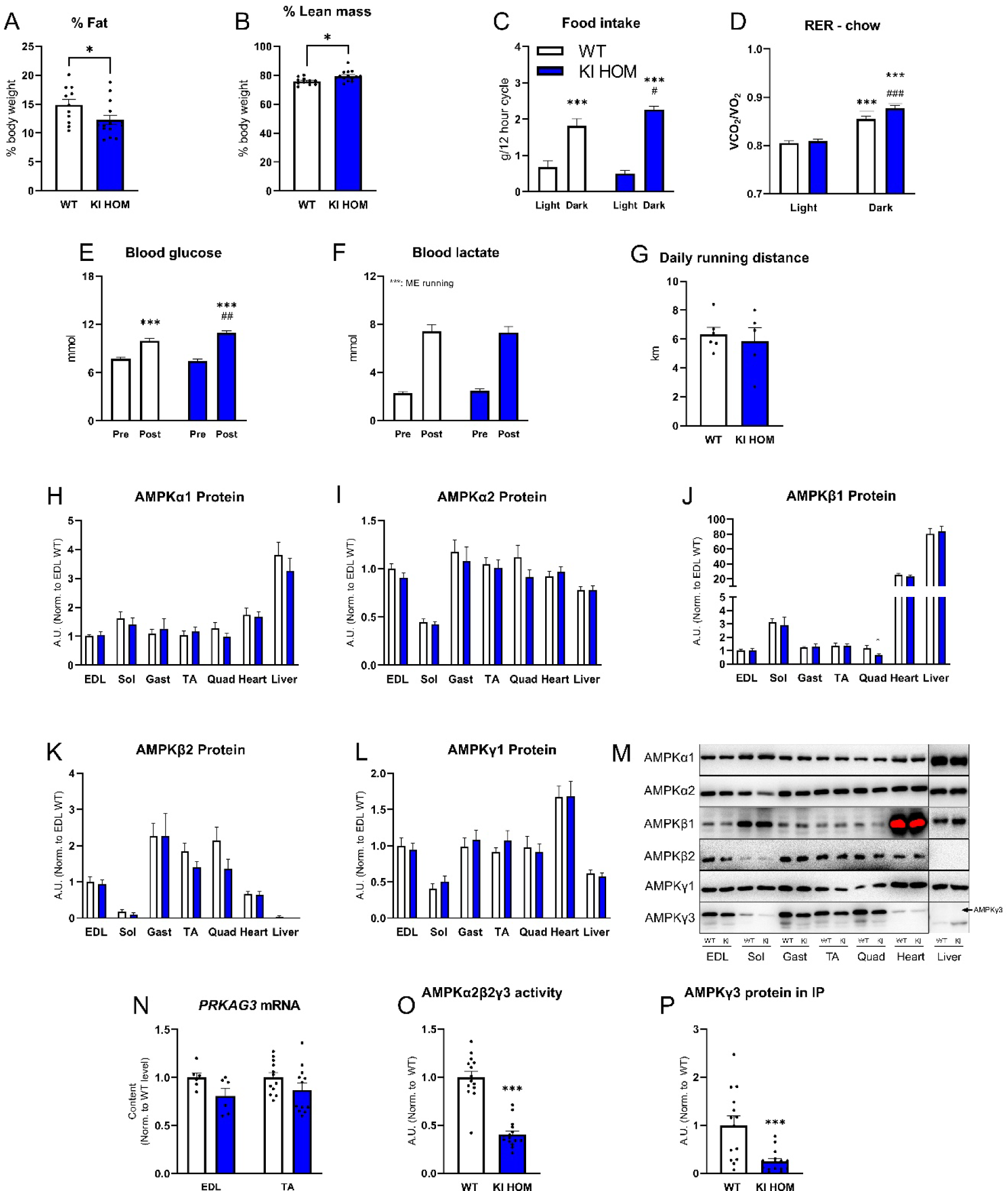
Characterization of KI HOM mice on a whole-body and muscle level. Wild-type (WT, white bar) and homozygote AMPKγ3 R225W carrier (KI HOM, blue bar) mice were investigated on a whole-body and muscle level. Amount of Fat (**A**) and lean mass (**B**) were related to total body weight (n=11-14). Food intake (**C**) and respiratory exchange ratio (RER) (**D**) were measured in metabolic chambers (n=11-13). Blood glucose (**E**) and blood lactate (**F**) concentrations were measured prior to and immediately after cessation of exhaustive treadmill running (n=8-9). Average daily voluntary running distance in running wheels (**G**, n=5-6). Protein expression of AMPK subunits α1 (**H**), α2 (**I**), β1 (**J**), β2 (**K**), γ1 (**L**) were measured in the indicated isolated muscles and tissues. Results are normalized to WT EDL levels for comparison (n=11). Representative Western blots (**M**). mRNA content of *PRKAG3* in EDL (**N**, n=6) and TA muscle (n=12). AMPKα2β2γ3 activity was measured in immunoprecipitations of AMPKγ3 protein from TA muscle in the basal and rested state (**O**, n=14-15). AMPKγ3 protein isolated in the AMPKγ3 immunoprecipitations used for the activity assay (**P**, n=14-15). Data are given as mean + SEM including univariate scatterplots to indicate individual values where applicable. Unpaired t-test (A-B, G, H-L, N-P) and Two-way ANOVA (C-F) were used to statistically evaluate differences between genotypes, running state/time of day or the interactions. *: Indicates difference between genotypes (A-B, G-P) or difference between running state/time of day within genotype (C-F). #: Indicates difference of genotype within running state/time of day (C-E). */#: p < 0.05, ##: p < 0.01, ***: p < 0.001. A.U.: Arbitrary Units, EDL: *m. extensor digitorum longus*, Sol: *m. soleus*, Gast: *m. gastrocnemius*, TA: *m. tibialis anterior*, Quad: *m. quadriceps*.

**Fig. S3.**
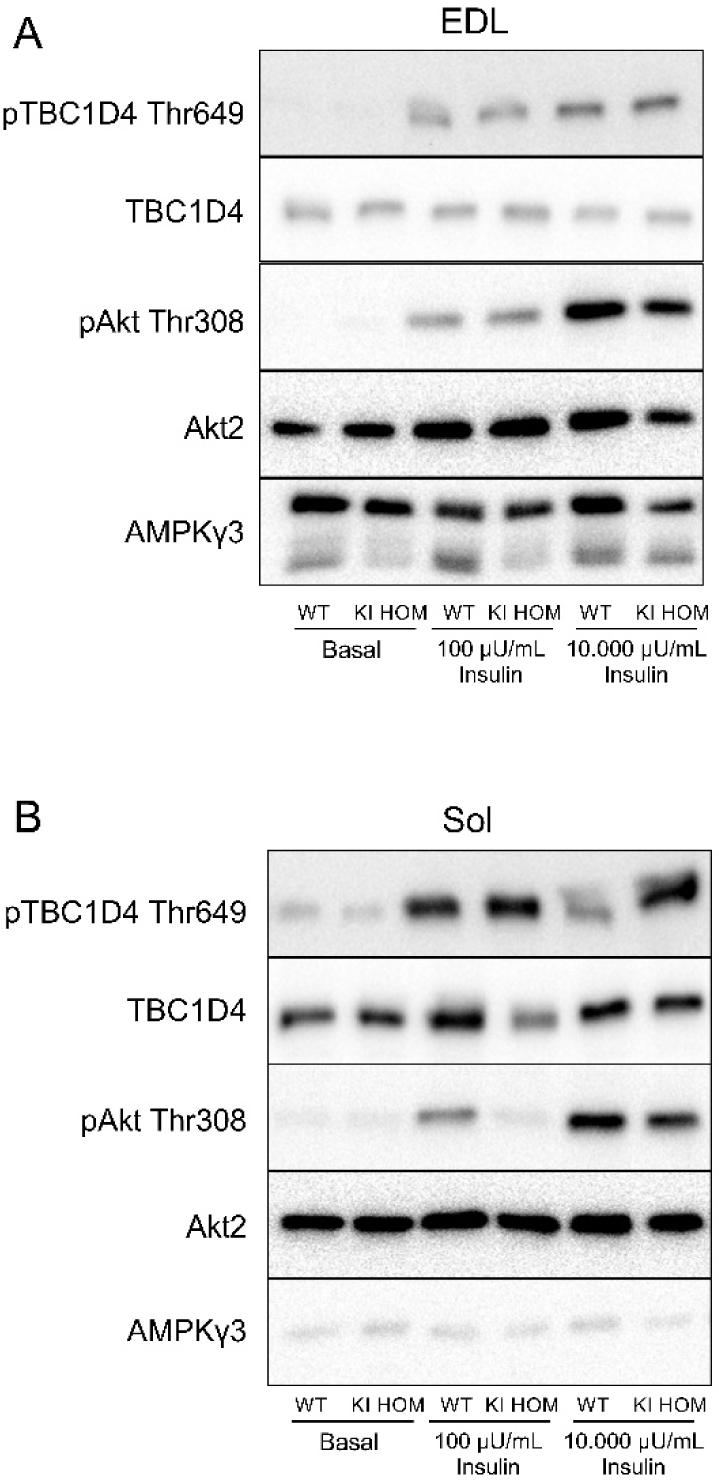
Representative Western blots during insulin stimulation in WT and KI HOM muscle. Muscle-specific insulin signaling was investigated in *m. extensor digitorum longus* (EDL) and *m. soleus* (Sol) from wild-type (WT) and homozygote AMPKγ3 R225W carrier (KI HOM) mice. Representative Western blots of pTBC1D4 Thr649, TBC1D4, pAkt Thr308, Akt2 and AMPKγ3 in EDL (**A**) and Sol (**B**) in the basal or insulin-stimulated state (100 µU/mL or 10.000 µU/mL).

**Fig. S4.**
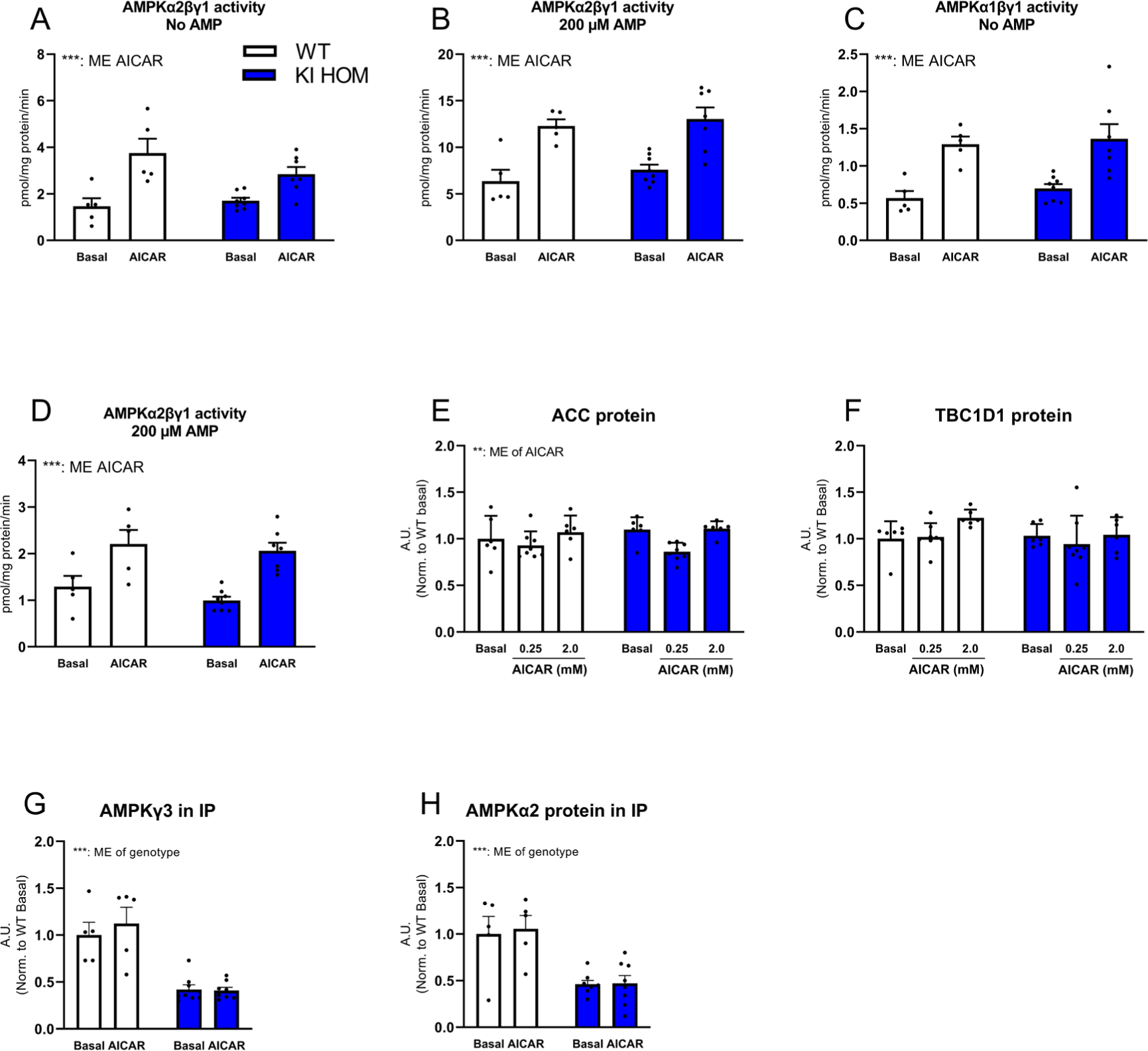
AMPK activity and signaling in AICAR-stimulated muscle from WT and KI HOM mice. To understand the function of the AMPKγ3 R225W protein, *m. extensor digitorum longus* (EDL) was isolated from wild-type (WT, white bar) and homozygote AMPKγ3 R225W carrier (KI HOM, blue bar) mice and stimulated with AICAR ex vivo. AMPKα2βγ1 (**A-B**) and AMPKα1βγ1 (**C-D**) activity was measured in the isolated EDL muscle in the basal or AICAR-stimulated state (2.0 mM) without or with 200 µM AMP added to the activity assay (n=5-8). Protein expression of ACC (**E**) and TBC1D1 (**F**) was measured in incubated EDL muscle in the basal state or with submaximal (0.25 mM) or maximal (2.0 mM) AICAR stimulation (n=6-8). The amount of AMPKγ3 (**G**) and AMPKα2 (**H**) in the AMPKγ3 immunoprecipitates. Data are given as means + SEM including univariate scatterplots to indicate individual values. All Western blotting data are normalized to WT basal levels. Two-way ANOVA (A-H) with Student-Newman-Keuls post-hoc test was used to evaluate differences between genotype, AICAR or the interaction. *: Indicates effect of AICAR compared to basal within genotype. #: Indicates difference between genotypes within AICAR concentration. **: p<0.01, ***/###: p<0.001. A.U.: Arbitrary Units. ME: Main Effect. IP: Immunoprecipitation.

**Fig. S5.**
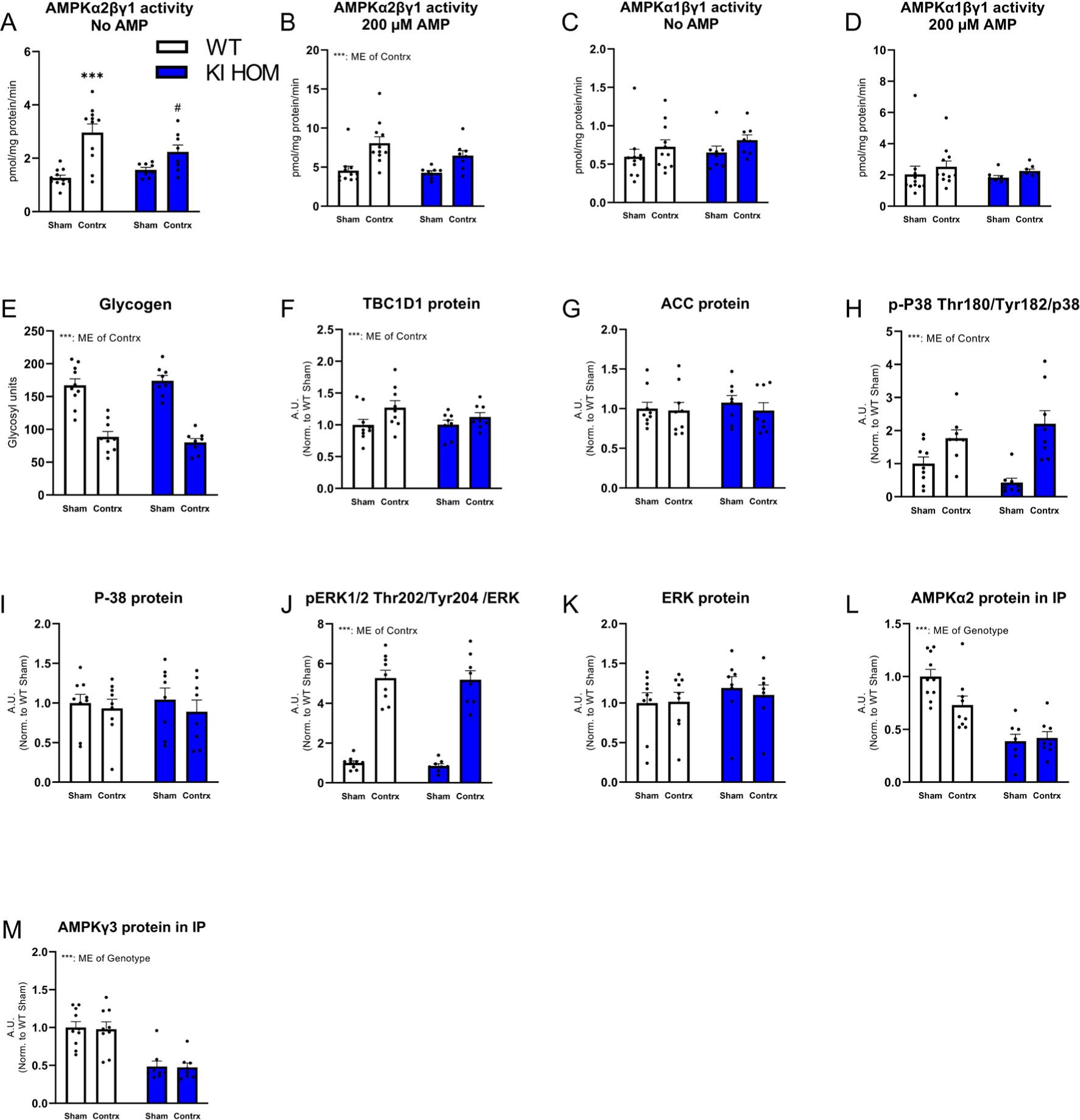
AMPK activity and signaling in contraction-stimulated muscle from WT and KI HOM mice. To understand the function of the AMPKγ3 R225W protein, m. tibialis anterior (TA) of a single lower hindlimb was contracted in wild-type (WT, white bar) and homozygote AMPKγ3 R225W carrier (KI HOM, blue bar) mice while the contralateral lower hindlimb served as a sham-operated resting control. AMPKα2βγ1 (**A-B**) and AMPKα1βγ1 (**C-D**) activity was measured in the sham-operated or contracted state without or with 200 µM AMP added to the activity assay (n=8-11). Glycogen levels in the sham and contracted muscles (**E**, n=8-10). Protein expression of TBC1D1 (**F**), ACC (**G**), p-P38 Thr180/Tyr182/P38 (**H**), p38 (**I**), pERK 1/2 Thr202/Tyr204/ERK (**J**) and ERK (**K**) was measured in sham or contracted TA (n=8-9). The amount of AMPKα2 (**L**) and AMPKγ3 (**M**) in the AMPKγ3 immunoprecipitates (n=8-10). Data are given as means + SEM including univariate scatterplots to indicate individual values. All Western blotting data are normalized to WT Sham levels. A two-way ANOVA (A-M) with Student-Newman-Keuls post-hoc test was used to evaluate differences between genotype, contraction or the interaction. *: Indicates effect of contraction compared to sham within genotype. #: Indicates difference between genotypes within contraction-state. #: p<0.05, ***/###: p<0.001. A.U.: Arbitrary Units. ME: Main Effect.

**Fig S6.**
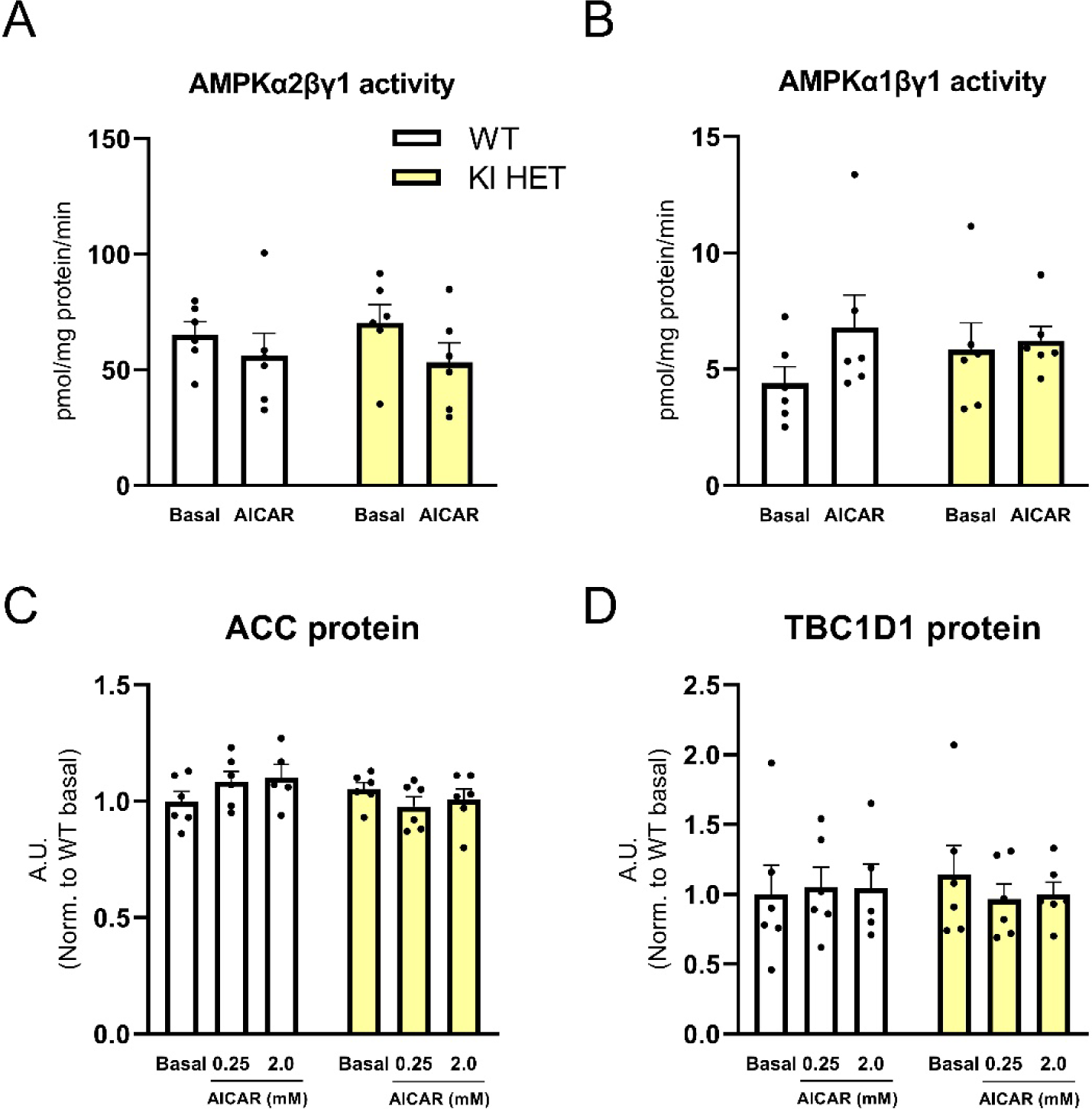
AMPK activity and signaling in AICAR-stimulated muscle from WT and KI HET mice. To introduce a combination of AMPKγ3 WT and AMPKγ3 R225W protein in skeletal muscle, heterozygote AMPKγ3 R225W carrier mice (KI HET, yellow bar) were generated and compared to wild-type mice (WT, white bar). AMPKα2βγ1 (**A**) and AMPKα1βγ1 (**B**) activity was measured in isolated EDL muscle in the basal or AICAR-stimulated state (2.0 mM) with 200 µM AMP added to the activity assay (n=6). Protein expression of ACC (**C**) and TBC1D1 (**D**) was measured in incubated EDL muscle in the basal state or under submaximal (0.25 mM) or maximal (2.0 mM) AICAR stimulation (n=5-6). Data are given as mean + SEM including univariate scatterplots to indicate individual values. A two-way ANOVA (A-D) with or without Student-Newman-Keuls post-hoc test was used to investigate differences between genotype, AICAR or the interaction. A.U.: Arbitrary Units. ME: Main Effect.

